# A freeze-and-thaw induced-fragment of the microtubule-associated protein Tau in rat brain extracts: implications for the biochemical assessment of neurotoxicity

**DOI:** 10.1101/404129

**Authors:** Israel C. Vasconcelos, Raquel M. Campos, Hanna K. Schwaemmle, Ana P. Masson, Gustavo D. Ferrari, Luciane C. Alberici, Vitor M. Faça, Norberto Garcia-Cairasco, Adriano Sebollela

**Affiliations:** Department of Biochemistry and Immunology, Ribeirao Preto Medical School, University of São Paulo, SP, 14049-900, Brazil; University of Tübingen, Germany; Department of Biomolecular Sciences, School of Pharmaceutical Sciences of Ribeirao Preto, University of São Paulo, SP, 14040-903, Brazil; Department of Physiology, Ribeirao Preto Medical School, University of São Paulo, SP, 14049-900, Brazil

**Keywords:** Tau protein, Neuropathology, Brain, Tissue homogenate, Proteolysis, Western blotting

## Abstract

Tau is a microtubule-associated protein responsible for controlling the stabilization of microtubules in neurons. Tau function is regulated by phosphorylation. However, in some neurological diseases Tau becomes aberrantly hyperphosphorylated, which contributes to the pathogenesis of neurological diseases, known as tauopathies. Western blotting (WB) has been widely employed to determine Tau levels in neurological disease models. However, Tau quantification by WB should be interpreted with care, as this approach has been recognized as prone to produce artifactual results if not properly performed. In this study, our goal was to evaluate the influence of a freeze-and-thaw cycle, a common procedure preceding WB, to the integrity of Tau in brain homogenates from rats, 3xTg-AD mice and human samples. Homogenates were prepared in ice-cold RIPA buffer supplemented with protease/phosphatase inhibitors. Immediately after centrifugation, an aliquot of the extracts was analyzed via WB to quantify total and phosphorylated Tau levels. The remaining aliquots were stored for at least 2 weeks at either −20°C or −80°C and then subjected to WB. Extracts from rodent brains submitted to freeze-and-thaw presented a ~25 kDa fragment immunoreactive to anti-Tau antibodies. An in-gel digestion followed by mass spectrometry analysis in excised bands revealed this ~25 kDa species corresponds to a Tau fragment. Freeze-and-thaw-induced Tau proteolysis was detected even when extracts were stored at −80°C. This phenomenon was not observed in human samples at any storage condition tested. Based on these findings, we strongly recommend the use of fresh extracts of brain samples in molecular analysis of Tau levels in rodents.

## INTRODUCTION

Tau is a neuronal, microtubule-associated protein (MAP) responsible for controlling the stabilization of microtubules in neurons, thereby impacting the coordination of the axoplasmic transport of organelles, proteins, lipids, synaptic vesicles and other important cargos along the neuron (1,2). Tau function is regulated by phosphorylation, but in some neurological diseases it is found aberrantly hyperphosphorylated (2,3). In fact, the involvement of key phosphorylation sites in Tau dysfunction, such as serine (Ser) 396, has been well demonstrated (4–6). Due to the strong correlation between high levels of hyperphosphorylated Tau, aggregation, microtubule destabilization and neuronal dysfunction/death, elevated hyperphosphorylated Tau levels have been assigned as a hallmark of several neuropathologies, collectively known as tauopathies (3,7–9). In addition, it has been shown that increased production of Tau protein fragments, due to enzymatic cleavage, also correlates with neurodegeneration in cellular models (10) and, more importantly, with memory deficits in animal models (11).

Western blotting (WB) has been widely employed in the assessment of Tau protein levels (either total or hyperphosphorylated Tau), as well as in the analysis of Tau fragmentation in a number of studies (11–14). However, quantification of Tau levels/cleavage by WB should be interpreted with care, as this approach has been recognized as a complex, multi-step technique that requires case-to-case standardization and is subjected to produce artifactual results if not performed properly (15,16). Of critical importance is the maintenance of the integrity of the protein of interest throughout processing, since unsuitable conditions for collection, storage, handling or experimental separation/detection may damage the target molecule. Thus, protein degradation could lead to conclusions that do not accurately reflect the biological phenomenon under investigation (15,17). This issue is particularly relevant in studies comprising the analysis of a large set of samples, such as in the case of studies using rodents, in which storage by freezing is common.

The goal of the present study was to evaluate the influence of freeze-and-thaw prior to WB analysis on the integrity of the Tau protein in extracts from rat brains. We found that rat brain extracts submitted to a single freeze-and-thaw cycle presented fragmentation of Tau protein, as detected by the presence of an extra ~25 kDa band simultaneously to the weakening of the bands corresponding to full-length Tau in samples prepared from both hippocampus and frontal cortex. Interestingly, Tau fragmentation was minor in brain extracts from 3xTg-AD mice, and absent in adult human brain extracts, suggesting that differences in Tau primary sequence between species may influence its liability to freeze and thaw-triggered degradation. Based on this finding, we recommend that molecular analyses of Tau levels/fragmentation in rat brain extracts should be carried out only using fresh extracts, in order to produce results that accurately reflect the biological phenomenon under investigation.

## METHODS

### Animals

The procedures were carried out using 12 month-old animals (*Rattus norvegicus)*, both control Wistar (n=7) and Wistar Audiogenic Rat (WAR) (n=9) strains (18), were approved by the Ethics Committee on Animal Use of Ribeirão Preto Medical School (CEUA-FMRP), under the protocol #017/2014-1. Control Wistar rats were obtained from the Central Vivarium of the University of São Paulo (USP) at Ribeirão Preto and kept at 25° C, 12h/12h photoperiod and water and food *ad libitum*. WAR animals were obtained from the Vivarium of the Department of Physiology of the USP Ribeirão Preto Medical School and kept at the same conditions. The procedures carried out using 6-8 months-old animals (*Mus musculus*) transgenic mice 3xTg-AD (n=6) strain (19), were approved by the Ethics Committee on Animal Use of School of Pharmaceutical Sciences of Ribeirao Preto (CEUA-FCFRP), under the protocol 18.1.6.60.1. The 3xTg-AD mice strain was obtained from the Vivarium I of the School of Pharmaceutical Sciences of Ribeirao Preto and kept at 23° C, 12h/12h photoperiod and water and food *ad libitum*.

### Ex-vivo human brain slices

The procedures carried out using *ex-vivo* human brain slices (20) were approved by the Ribeirão Preto Medical School Ethics Committee, under the protocol HCRP #17578/15. Briefly, a fragment of human frontal cortex was surgically collected, sliced (200μm thick) and cultured for 2 days, as described in detail in (20,21).

### Western Blotting

Animals were anaesthetized by inhalation of isoflurane (*BioChimico*, Brazil) and euthanized by decapitation. Cerebral tissue was collected, dissected and stored in 1.5 mL microtubes at −80 °C. Slices were stored at 1,5 mL microtubes immediately prior to homogenization. Homogenates from dorsal portion of hippocampus (rat), frontal cortex (mice and rat) and *ex-vivo* human brain slices were prepared according to Petry et al. (22). Tissue fragments were homogenized in ice-cold RIPA buffer (50 mM Tris, 150 mM NaCl, 1 mM EDTA, 1 % Triton X-100 and 0.1 % SDS, pH 7.5) supplemented with protease inhibitor (1:100, Sigma) and phosphatase inhibitor (10 mM NaF, 10 mM Na_3_VO_4_) cocktails at the ratio of 10 μL per mg of tissue using an electric potter (Kimble Chase) in 3 cycles of 10 seconds each. Extracts were centrifuged for 10 minutes at 10000 x g and 4 °C. Supernatant was transferred to new microtubes and aliquots were readily used for analysis of Tau protein by WB (fresh extract). Alternatively, parts of the extracts were stored at −20 °C and −80 °C for at least 2 weeks and then thawed just before WB (freeze/thaw extract). Preparation of samples for SDS-PAGE included addition of protein loading buffer (0.0625 M Tris-Cl, 2 % SDS, 10 % v/v glycerol, 0.1 M dithiothreitol, 0.01 % bromophenol blue, pH 6.8) to extracts and boiling for 5 minutes at 100 °C. The SDS-PAGE was carried out using 12 % acrylamide/bis-acrylamide gels and a constant voltage of 90 V. For blotting, 0.45 μm nitrocellulose membranes (GE *Healthcare Life Sciences*) were utilized. Membranes were blocked in 5 % nonfat dry milk/T-TBS (Tris-buffered saline: 20 mM Tris pH 7,5, 150 nM NaCl, 0,1 % Tween 20) solution for 1 h at RT and subsequently incubated with primary antibody overnight at 4 °C. Primary antibodies used for Tau assessment were anti-Tau phospho S396 (*Abcam*) or anti-Tau [E178] (*Abcam*), both 1:1000 in 5 % BSA/T-TBS solution. After washing, membranes were incubated with secondary antibody ECL anti-rabbit IgG HRP (*Amersham*) at 1:3000 in 5 % BSA/T-TBS for 1 h at RT. The membranes were revealed using *ECL Prime Western Blotting Detection Reagent* (*GE Healthcare Life Sciences*) and imaged in *ChemiDoc* imaging system (*Bio-Rad*) coupled to a digital system and software *ImageQuant*^TM^ 3.5 (*GE Healthcare Life Sciences*). For β-actin probing, membranes were initially deblotted with stripping solution (10% SDS, 100 mM glicin, 0,1% NP40, pH 2,5) for 30 minutes under vigorous shaking, washed in T-TBS solution then incubated with anti-β-actin (*EMD Millipore*) at 1:20000 in 3 % BSA/T-TBS for 1 h at RT.

### In-gel digestion and LC-MS/MS analysis

Aliquots (250 μg total protein) of hippocampal extracts of WAR animals subjected to freeze/thaw were treated with 5 μL of dithiothreitol (*Biorad*) 50 μg/μL, diluted in ammonium bicarbonate (*Sigma*) 100 mM, pH 7.8, for 30 min at 37 °C. DTT-treated samples were then diluted in loading buffer, boiled for 5 min at 100 °C, and alkylated in the dark for 30 min at 37 °C with 1250 μg of iodoacetamide (*Sigma*) 50 μg/μL, diluted in ammonium bicarbonate 100 mM, pH 7.8. After sample separation by SDS-PAGE (carried out using 12% acrylamide/bis-acrylamide gels and a constant voltage of 90 V), a piece of ca. 30 mm^2^ of the unstained gel (per lane) was cut off using as a reference the 25 kDa standard band (pre-stained). Gel pieces corresponding to the fragment were further sliced to 1-4 mm^2^ pieces, washed and digested with trypsin (*Promega*) as described by Grassi et al. (2017). Tryptic peptides were successively extracted with 5 % formic acid (*Merck*)/50 % acetonitrile (*Merck*), and then 90 % acetonitrile, and dried in a vacuum concentrator. The sample was re-suspended in 50 μL 50 % acetonitrile/5 % formic acid and centrifuged at 12000 xg for 15 min at 25 °C. The supernatant was injected into a LC-MS/MS *Xevo TQS system* (*Waters*). Chromatographic separation was performed in a UPLC (I-class, *Waters*) using a C18 column (1.8 μm particle size, 100 Å pore size, 1 mm × 150 mm, Waters) in a linear gradient of 5 to 30 % acetonitrile over 15 min at 100 μL/min in a formic acid:acetonitrile:water solvent system. Detection of Tau theoretical tryptic peptides was programmed in a scheduled multiple reaction monitoring method using 3–5 transitions per peptide and monitoring windows of 2 min centered on predicted retention time. Both method development and data analysis were conducted using *Skyline* software (23).

### Sequences alignment

The Tau sequences for rat (P19332), mouse (P10637) and human (P10636) were obtained from the Uniprot databank (https://www.uniprot.org/) and analyzed using AliView viewer and editing tool (24).

## RESULTS

### Presence of an anti-Tau-immunoreactive ~25 kDa band in rat brain extracts submitted to freeze-and-thaw

Elevated Tau phosphorylation levels, particularly in key phosphorylation sites such as Ser^396^, are known to be associated with cognitive impairment and the pathogenesis of several neuropathologies (2,4–7). With the purpose of investigating a possible involvement of Tau hyperphosphorylation in a cohort of animals presenting early cognitive decline (25,26), we initially prepared hippocampal extracts from 12-month old Wistar rats and evaluated Tau protein levels, both phosphorylated at Ser^388^ (analogue to Ser^396^ in humans) and total Tau. In parallel, we determined both total and phospho-Ser^388^ Tau levels in extracts from 12-month old WAR animals – a rat strain used as a model of epilepsy and neuropsychiatric comorbidities such as depression and anxiety (27). The brain extracts had been previously prepared and stored at −20 °C. Unexpectedly, in addition to the typical profile of three major bands migrating between 50 and 75 kDa (Fig. 1A), which are known to correspond to different isoforms of Tau expressed in rodent brains (28), we found an extra band of ~25 kDa in all samples probed with either anti-phospho or total Tau antibodies. The ~25 kDa band was also observed in extracts from frontal cortex stored and handled likewise (Fig. 1B).

**Figure 1.**
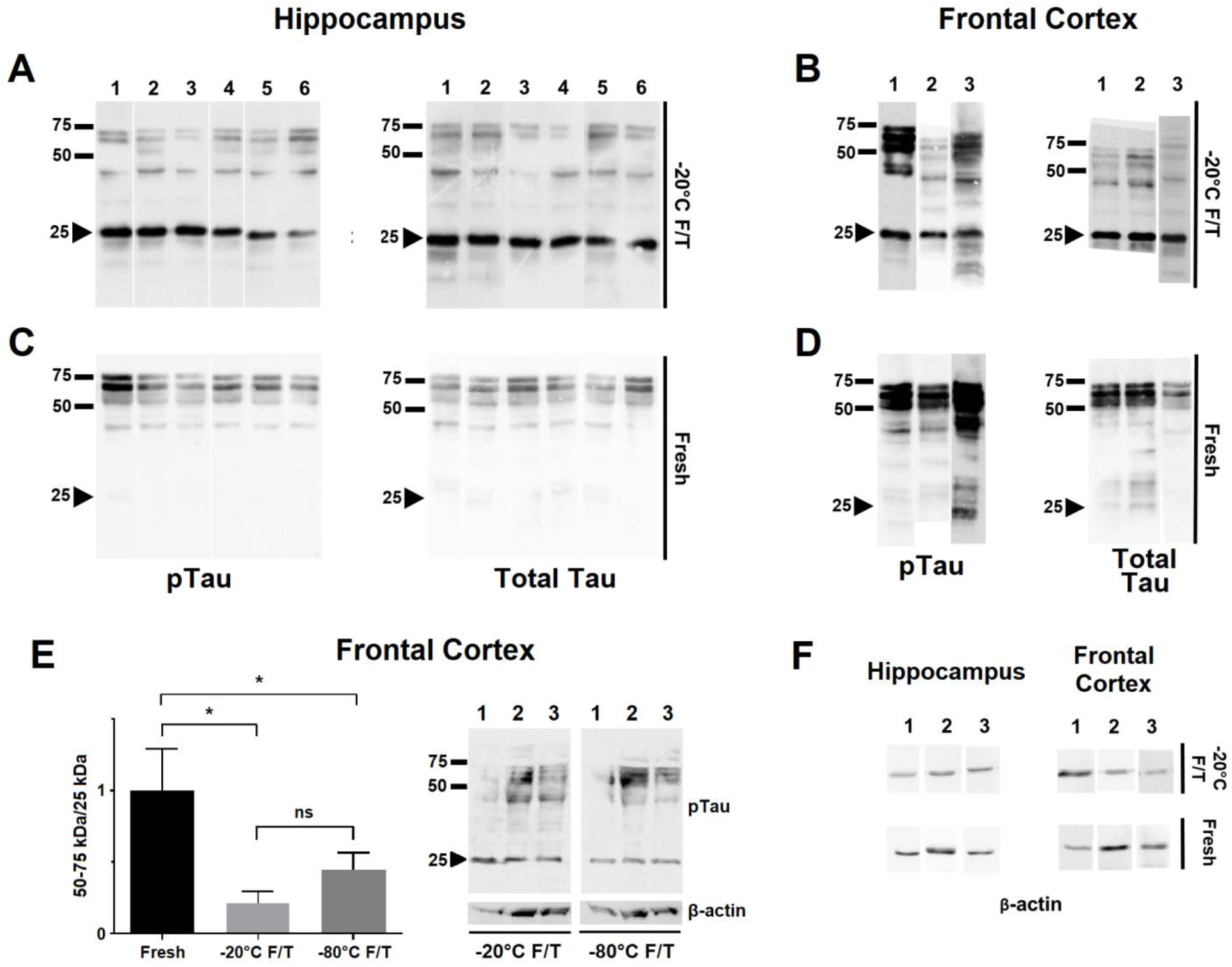
Presence of a ~25 kDa anti-Tau immunoreactive band in freeze-and-thawed brain extracts of 12-month old Wistar rats. Extracts were analyzed by Western blotting after a freeze-and-thaw cycle at −20 °C (−20 °C F/T; A and B) or immediately after preparation (fresh; C and D). Membranes were probed with either anti-Tau phospho S396 (pTau; A-D and F) or anti-Tau [E178] (Total Tau; A-D) antibodies. Extracts were prepared from dorsal hippocampus (A and C) or frontal cortex (B and D). (E) pTau/25kDa band ratio. A representative WB comparing extracts from frontal cortex stored at different temperatures is shown (n = 3 per group; *p<0.05). Some extracts were additionally probed for β-actin (F). The arrow-head highlights the ~25 kDa regions on the membranes. Some samples (A: lanes 1-3, 9, 10 and 12; B: lanes 2-5; C: lanes 3 and 6) were obtained from WAR animals, all the remaining samples were obtained from wild-type Wistar rats.

Zhao et al. (11) have reported the presence of a ~35 kDa (TCP35) band in transgenic mice brains expressing human Tau, which was correlated with cognitive decline and identified as a proteolytic product of Tau. Motivated by those findings, and considering that freeze-and-thawing may impact the stability of several proteins (29–32), we wondered whether this ~25 kDa fragment observed under our conditions was endogenously produced or rather corresponded to a product of freeze-and-thaw-induced degradation. Interestingly, when the Western blotting was conducted using fresh samples, the ~25 kDa band was significantly weaker or even completely absent in both hippocampal (Fig. 1C) and frontal cortex (Fig. 1D) extracts. Importantly, the bands corresponding to full-length Tau were markedly weaker when the ~25 kDa band was present, supporting the notion that it represents a degradation product of Tau generated by freeze-and-thawing the extracts.

Next, we investigated whether storing the extracts at −80 °C, instead of −20 °C, would prevent the appearance of the ~25 kDa band and/or weakening of full-length Tau bands. Data obtained using a different group of animals indicated that storage at −80 °C did not prevent the freeze-and-thaw induced phenomenon, since presence of the ~25 kDa band and the weakening of 50-75 kDa bands, and consequently the ratio full-length Tau/25 kDa fragment was similar to that of samples stored at −20 °C (Fig. 1E). Furthermore, this freeze and thaw-induced phenomenon does not seem to equally affect different proteins in the extracts, since no differences in integrity of β-actin between fresh and freeze/thaw extracts were observed (Fig. 1E and 1F). The appearance of the ~25 kDa fragment, which originated from freeze-and-thawing the brain extracts, was detected at similar degrees in extracts of both Wistar and WAR strains, indicating that this phenomenon is rat strain-independent.

### Identification of the ~25 kDa band associated to freeze-and-thaw as a degradation product of Tau

In order to verify whether freeze-and-thawing caused Tau fragmentation, we aimed to biochemically identify the freeze-and-thaw-induced ~25 kDa species by mass spectrometry (MS). For this purpose, an extract of hippocampus from a WAR rat stored at −20 °C was thawed and submitted to protein separation by SDS-PAGE. A piece of the gel in the range of 25 kDa was cut off and subjected to trypsin digestion. The eluted peptides were analyzed by target protein MS. Nine peptides of full-length rat Tau sequence were identified, one of those being a unique peptide (Table 1). This result confirmed that the ~25 kDa band showing immunoreactivity to anti-Tau antibodies is indeed a degradation product of Tau. All of the Tau-associated peptides identified were located near the C-terminus (Fig. 2).

**Table I.**
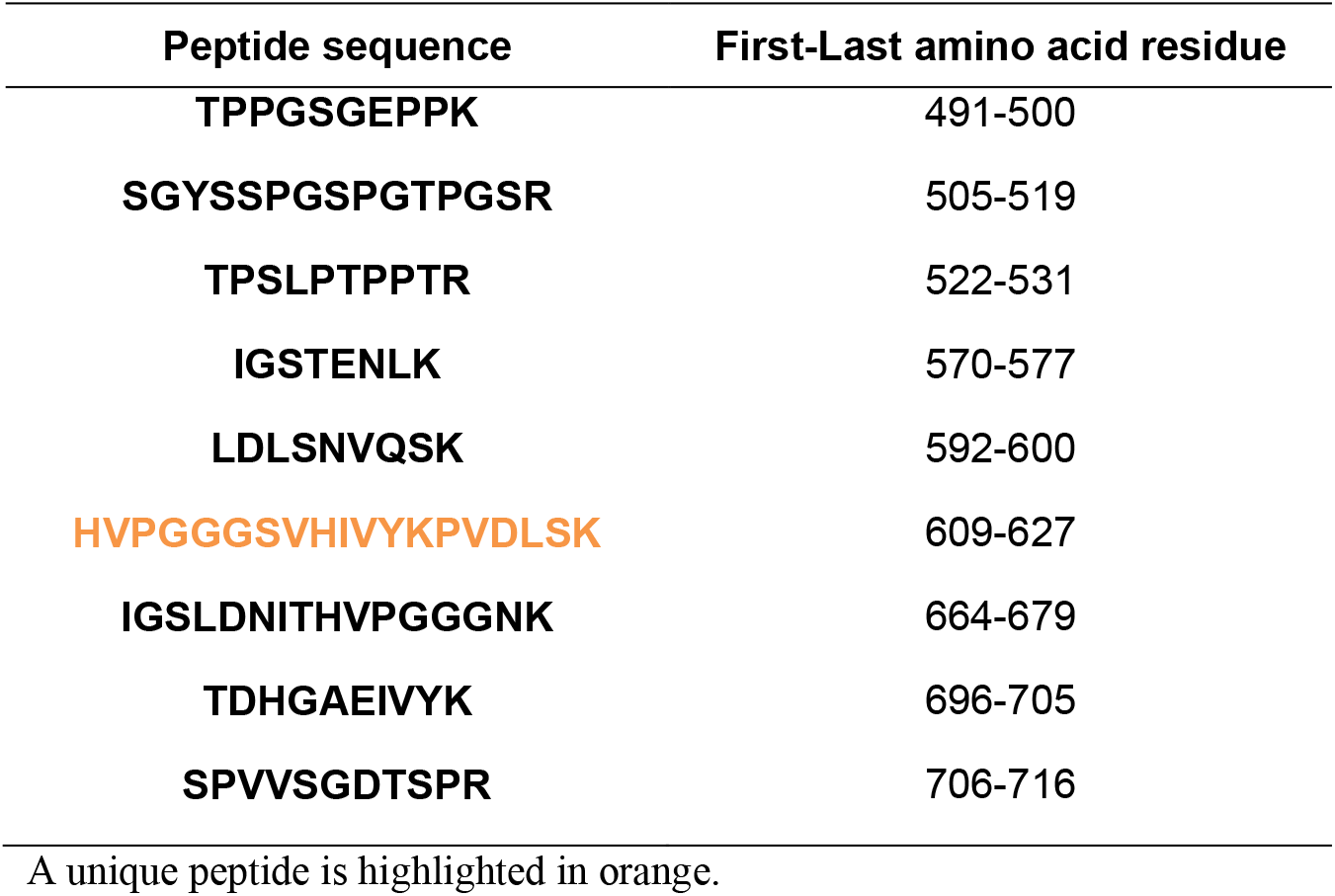
Tryptic peptides identified by targeted mass spectrometry.

**Figure 2.**
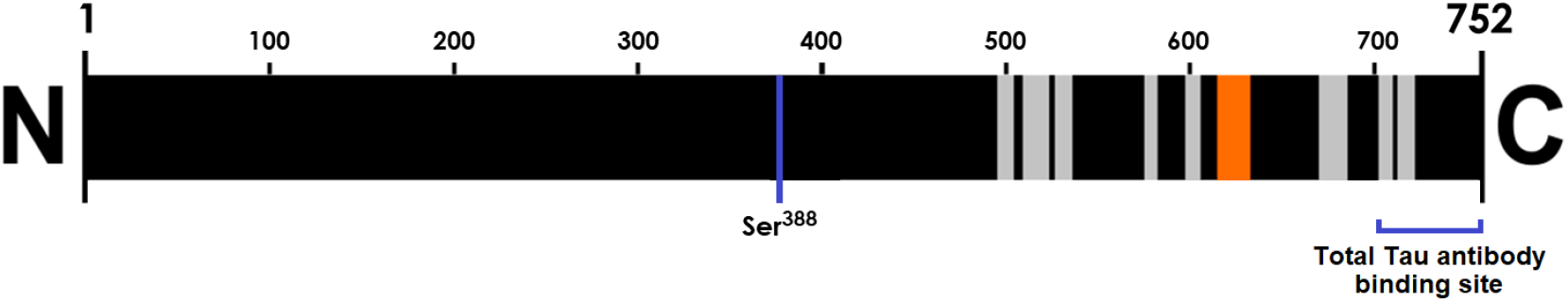
Localization of Tau-derived peptides identified by mass spectrometry in full length Tau sequence. All the peptides identified in the mass spectrometry fragment analysis are shown in gray (a unique peptide is highlighted in orange). All the tryptic peptides are located after the Ser388 (human analogue of human Ser^396^) and near to the C-terminus of full-length Tau from rat. The total Tau antibody binding site, indicated in the C-terminus, refers to the region where the proprietary sequence is located.

Considering the canonical sequence of rat Tau protein (uniprot.org/uniprot/P19332), and the immunoreactivity of the ~25 kDa fragment to anti-Tau phospho S396 antibody (the corresponding residue to human Tau Ser^396^ in rats being Ser^388^), we mapped the cleavage site associated to the generation of the 25kDa fragment to be in between the N-terminus and the Ser^388^. Moreover, the anti-total Tau E178 antibody used in the WB analyses also reassures the truncated portion contains the C-terminus of Tau protein.

### Differences in the generation of the ~25 kDa Tau fragment in samples from a mice brain disease model and human samples

Tau dysregulation is largely involved in neurodegenerative processes (3,8,9), and more recently, Tau fragmentation has been associated to cognitive decline in mice models of Alzheimer’s disease (11). After identifying the 25 kDa species as a degradation product of Tau, we wondered whether this phenomenon observed in rat brain samples, including the seizure-prone WAR rat strain, would also take place in samples from a different rodent experimental model of neuropathology. For this purpose, we used frontal cortex extracts from 3xTg-AD mice – a widely used animal model of Alzheimer disease (19). Similarly to the observed 25 kDa species in samples from rats, the extracts from 3xTg mice brain presented Tau degradation, i.e. the appearance of a ~25 kDa fragment along with a slight reduction in bands corresponding to full-length Tau (Fig. 3A), which led to a 42 % reduction in the ratio between full-length Tau/25 kDa fragment levels (Fig. 3B). However, in this cohort this reduction reached no statistical significance.

**Figure 3.**
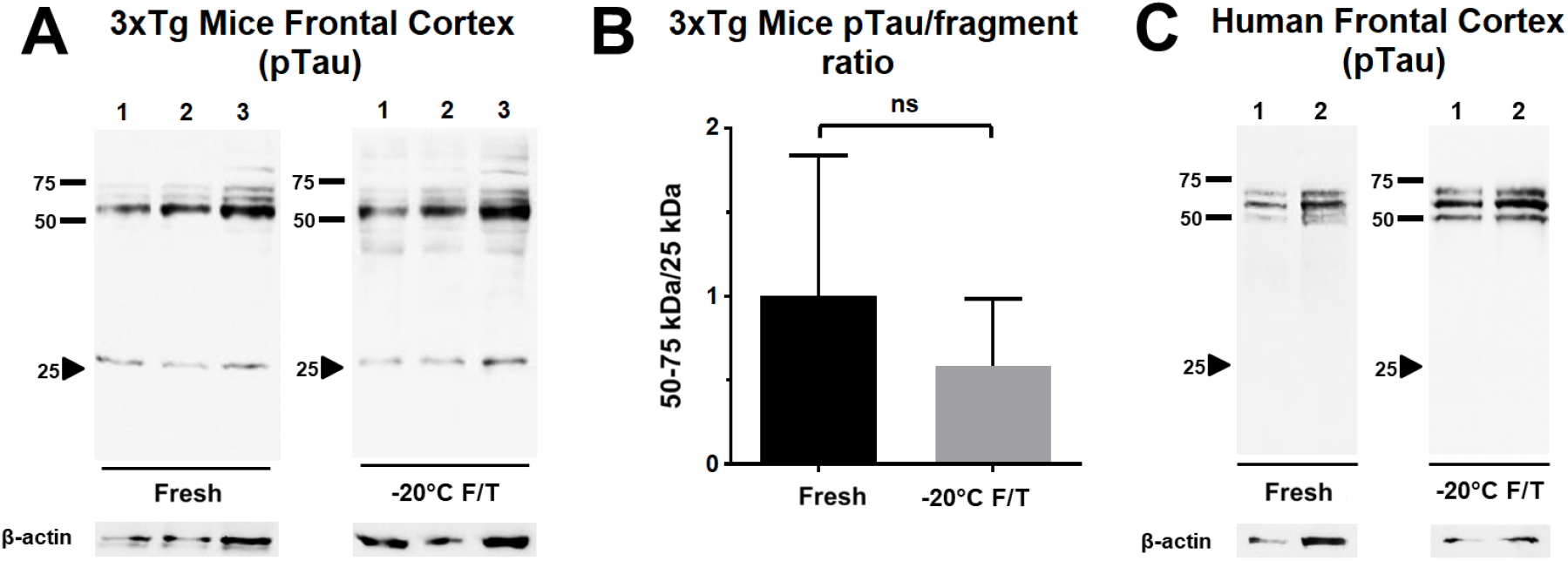
Evaluation of freeze and thaw-induced Tau degradation in extracts from experimental models of neurodegeneration. (A) Representative WB images of 3xTg mice frontal cortex extracts probed for pTau and β-actin. Extracts were either analyzed fresh or after storage at (−20 ºC F/T). The arrow-head indicates the presence of a band near 25 kDa in both fresh and −20°C F/T. (B) Full length/fragment ratio obtained from the quantification of 3xTg frontal cortex WB analysis (n = 6 for each group; *t* test *p<0.05). [C] The same WB analysis but using human brain slices extracts obtained from two donors.

We also tested a non-rodent experimental model, slice cultures from adult human brains (20). Surprisingly, the ~25 kDa fragment was not observed in any condition (Fig. 3C), suggesting no apparent freeze and thaw induced Tau degradation takes places in human samples. A possible explanation for the differences observed in Tau degradation between rats, transgenic mice and human samples could be the differences in the primary sequences of Tau between the species, with likely implications to 3D structure, and thus to proteolysis propensity.

In order to estimate the possible contribution of amino acid residue differences to Tau stability in the Tau primary sequence between species, we performed a comparison of the canonical sequences of Tau from rat, mouse and human (Uniprot code: P19332, P10637 and P10636, respectively) using AliView (24). The sequence alignment comparison exhibits highly conserved sequences around the serine residue targeted by the anti-Tau phospho S^396^ antibody (Fig. 4). Despite the similarities, three non-conservative changes are present regarding rodents *versus* human sequences: an arginine (R^377^_rat_; R^357^_mouse_) residue is substituted by a glycine (G^385^_human_), a proline (P^390^_rat_; P^371^_mouse_) is substituted by an arginine (R^398^_human_), a proline (P^396^_rat_; P^377^_mouse_) is substituted by a leucine (L^404^_human_) and a serine (S^397^_rat_; S^378^_mouse_) residue is substituted by a lysine (K^405^_human_). Taking the substantial differences regarding the charge, polarity or structure flexibility assigned by those residues into account, it is possible to infer that those differences may play a key role in the proteolysis propensity seen in rodent Tau proteins.

**Figure 4.**
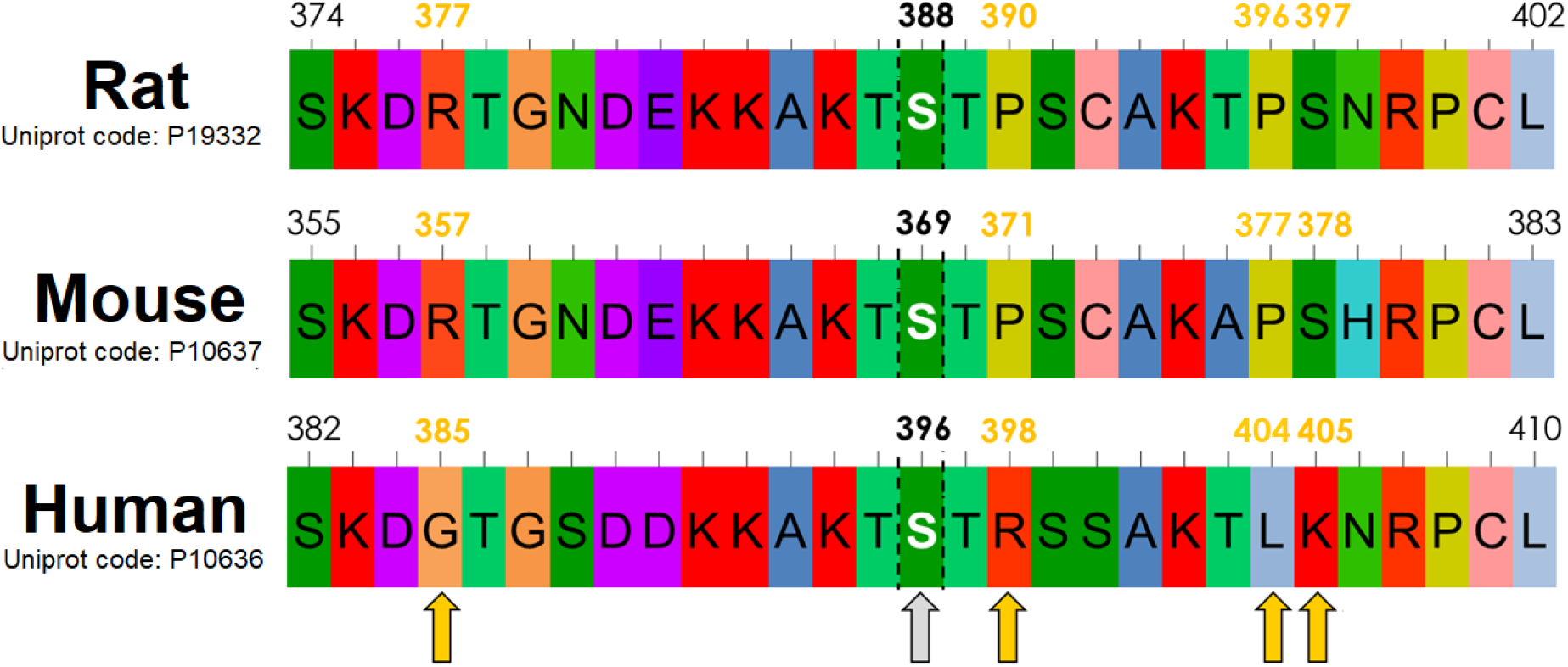
Sequence comparison of Tau residues around phospho Ser^396^ epitope in rodents and humans. Comparison between Tau sequences from rat, mouse and human around the serine-phosphorylation site (gray arrow) probed by the anti-Tau phospho S396 antibody (Ser^396^ analogue in rat is Ser^388^; in mouse is S^369^). Considering the differences in reactivity and biochemical nature of side chains, the positions where significant substitutions are present comparing rodent *versus* human sequences are indicated by yellow arrows. The default color scheme of AliView was used in the alignment.

## DISCUSSION

In this work we present results indicating that a ~25 kDa fragment generated by Tau degradation is present when extracts from rat brains are subjected to freeze and thaw – a common step prior to WB analyzes. Although reports of protein degradation in brain samples stored as frozen extracts have been shown for samples stored in the absence of protease inhibitors (33), our present study shows the occurrence of proteolysis even in extracts prepared using a protease inhibitor cocktail. Surprisingly, this phenomenon was minor in 3xTg mice brains, an animal model of neurodegeneration, and absent in samples from human brains, suggesting that differences in primary sequences of Tau influence the liability of this protein to freeze and thaw induced proteolysis.

Given the widespread use of WB to determine Tau levels in brain extracts in experimental models of neurodegeneration, it is reasonable to raise a concern on the possible impact of freeze and thaw induced Tau proteolysis on the reliability of reported findings in which rodent Tau levels have been determined by WB. The emergence of the 25 kDa band, concomitant to the weakening of the bands corresponding to full length Tau (which are used to quantify the levels of Tau) might therefore introduce an artifactual outcome.

Assuming the freeze and thaw induced cleavage occurs near Ser^388^ (the analogue residue of human Ser^396^ in Tau sequence from rats), the predicted molecular weight of the C-terminal fragment is ~35 kDa. Although the reason for the ~10 kDa discrepancy between the apparent MW we have observed experimentally and the predicted molecular weight is not clear, it is well known that several proteins present anomalous behavior in MW determination by SDS-PAGE (e.g. (34)). It is also important to consider that Tau mRNA alternative splicing generates several isoforms of rat Tau (uniprot.org/uniprot/P19332#cross_references), and this may implicate in alternative Tau isoforms with unpredicted MW (35). Further analyses are necessary in order to determine which isoform has been cleaved during freeze-and-thaw to generate the ~25 kDa fragment detected in our experiments. Interestingly, the alleged cleavage region hypothesized in this study is close to a proteolysis site for caspase, mapped in human Tau isoform 0N4R (amino acid sequence: HVLGGGSVQIVYKPVD) by Zhao et al. (11). It is thus possible that this region, which is partially conserved and present in all isoforms of rat Tau (uniprot.org/uniprot/P19332), represents a proteolysis-prone site. Moreover, the conformational flexibility of the region surrounding residue Ser^396^ of human Tau has been recently demonstrated by the Ser^396^ phosphorylation-induced α-helix to β-sheet conversion, with possible implications to microtubule binding and aggregation (36).

The importance of these findings is reinforced when we take into account that the freeze and thaw induced Tau proteolysis was observed in samples from both wild-type (Wistar) and WAR rats, the later a strain that has been used to model epilepsy and associated comorbidities (27). Importantly, WAR animals have also recently shown to present spatial memory and learning deficits that correlate with increased Tau phosphorylation in hippocampus (25,26,37). Moreover, phosphorylated Tau levels are widely used as an indicator of neurodegeneration, in particular the phosphorylated epitope evaluated in this work, (Ser^396^ in human sequence), which is implicated in the pathophysiology of several neurological diseases such as Alzheimer’s and Pick disease (2–6). Therefore, it is not possible to rule out that, at least in some studies analyzing rodent Tau levels by Western blotting (e.g. 38–40), the reported quantification may have been impacted by the sample storage process, notably freeze and thawing brain extracts. Unfortunately, in many studies reporting Tau level analysis by WB, no detailed description of sample handling prior to WB is depicted. In addition, the representative images presented in many works (e.g. 41–45) usually do not allow a comprehensive evaluation on the presence of Tau fragments, due to membrane image cutting.

In summary, we report that freeze-and-thawing rat brain extracts led to Tau protein degradation, detected by the presence of an extra band of ~25 kDa, simultaneously to weakening of the three major bands corresponding to full-length Tau in WB analysis. Based on this observation, we strongly recommend performing molecular analysis of Tau levels in rodent brain samples using fresh extracts. Considering the wide use of WB to evaluate neurodegeneration in tauopathies, we anticipate that following this recommendation will lead to more accurate interpretations in future studies on Tau-related disorders.

## DATA AVAILABILITY

The authors confirm that the data supporting the findings of this study are available within the article and its supplementary materials. The raw data underlying the WB results reported are presented in the Supplementary Material.

## AUTHOR CONTRIBUTIONS

Vasconcelos, I. C., Campos, R. M.; Designed and performed experiments, analyzed data and co-wrote the paper. Schwaemmle, H. K.; Performed WB experiments for rat samples. Masson, A. P., Faça, V. M.; Designed and Performed MS experiments. Ferrari, G. D.; Performed dissection and tissue harvesting for 3xTg mice. Alberici, L. C.; Provided 3xTg mouse strain. Garcia-Cairasco, N.; Provided Wistar and WAR rat strain. Sebollela, A.; Designed experiments and co-wrote the paper.

## ACKNOWLEDGEMENTS

We thank Jose Antonio C. de Oliveira and Silvana El-Chedraoui Silva for excellent technical assistance.

