## Supplementary Material for "A freeze-and-thaw induced-fragment of the microtubule-associated protein Tau in rat brain extracts: implications for the biochemical assessment of neurotoxicity"

**Figure S1.** Raw data of **Fig.1 – Panel A** western blotting of -20°C freeze/thawed rat hippocampus extracts (lanes used in the main figure are indicated by dotted frames).

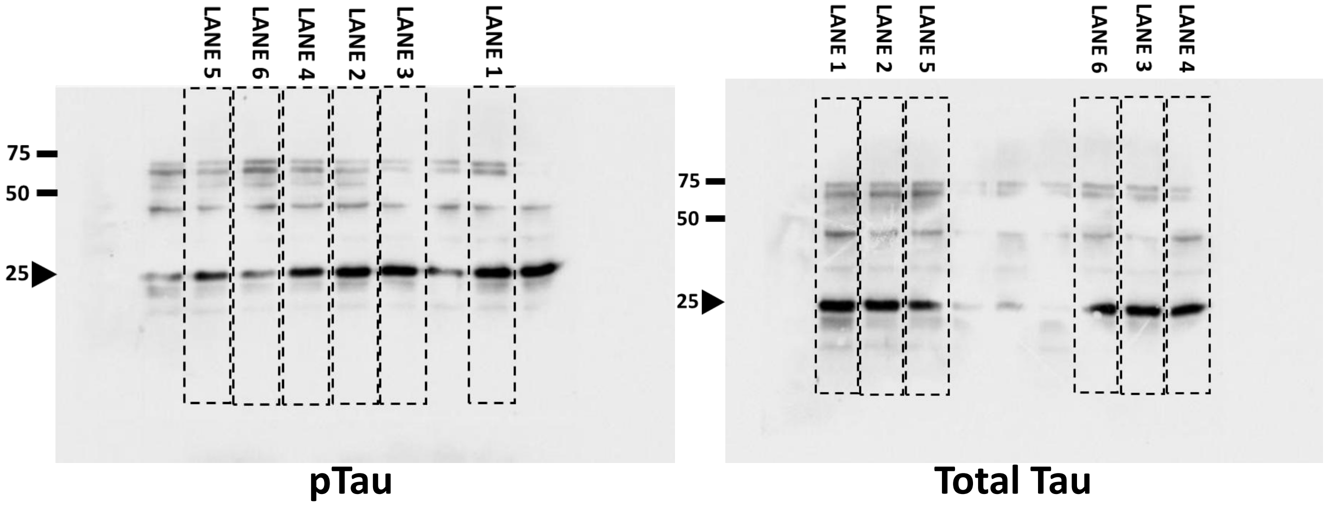

**Figure S2.** Raw data of **Fig.1 – Panel B** western blotting of -20°C freeze/thawed frontal rat cortex extracts (lanes used in the main figure are indicated by dotted frames).

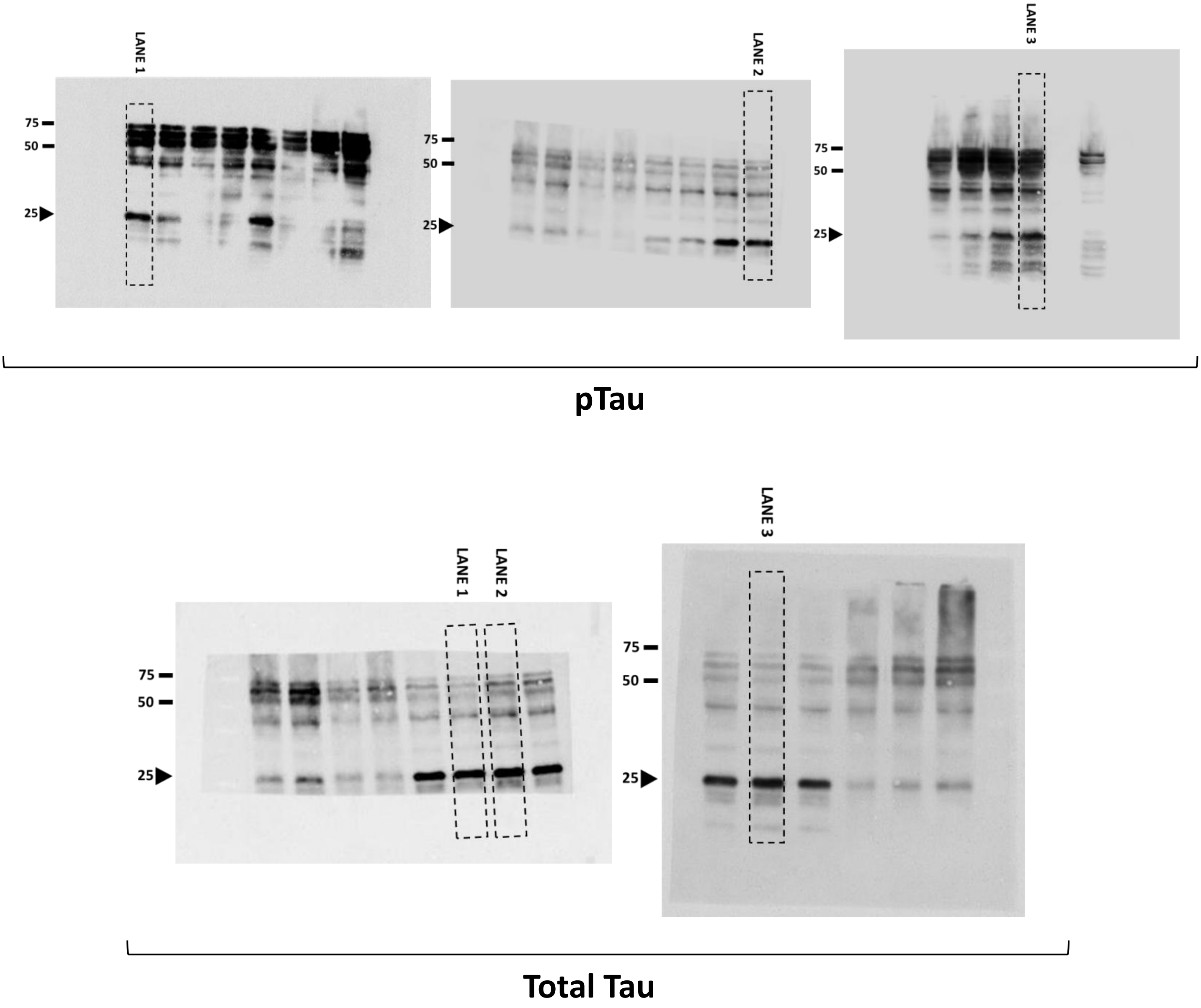

**Figure S3.** Raw data of **Fig.1 – Panel C** western blotting of fresh rat hippocampus extracts (lanes used in the main figure are indicated by dotted frames).

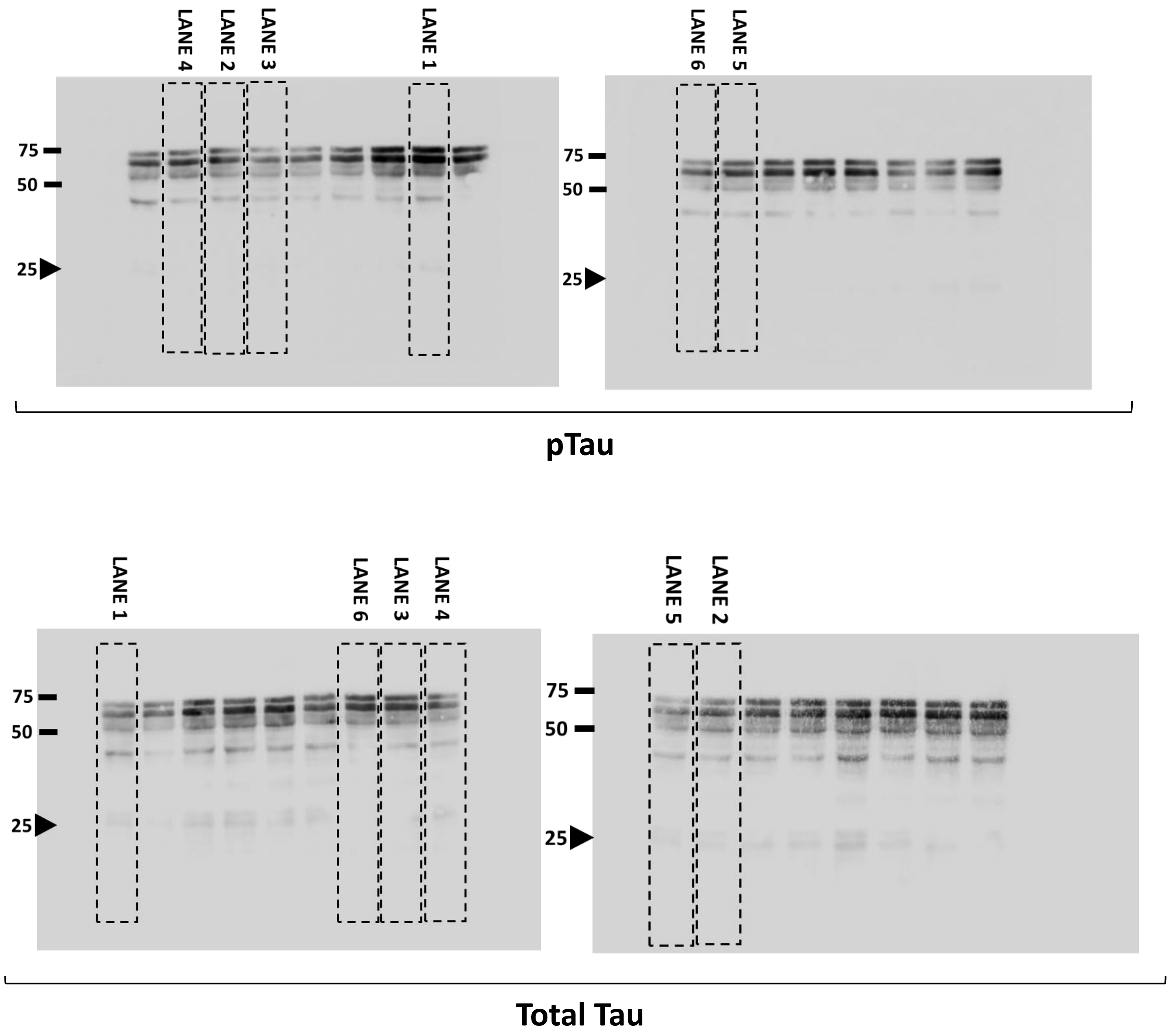

**Figure S4.** Raw data of **Fig.1 – Panel D** western blotting of fresh rat frontal cortex extracts (lanes used in the main figure are indicated by dotted frames).

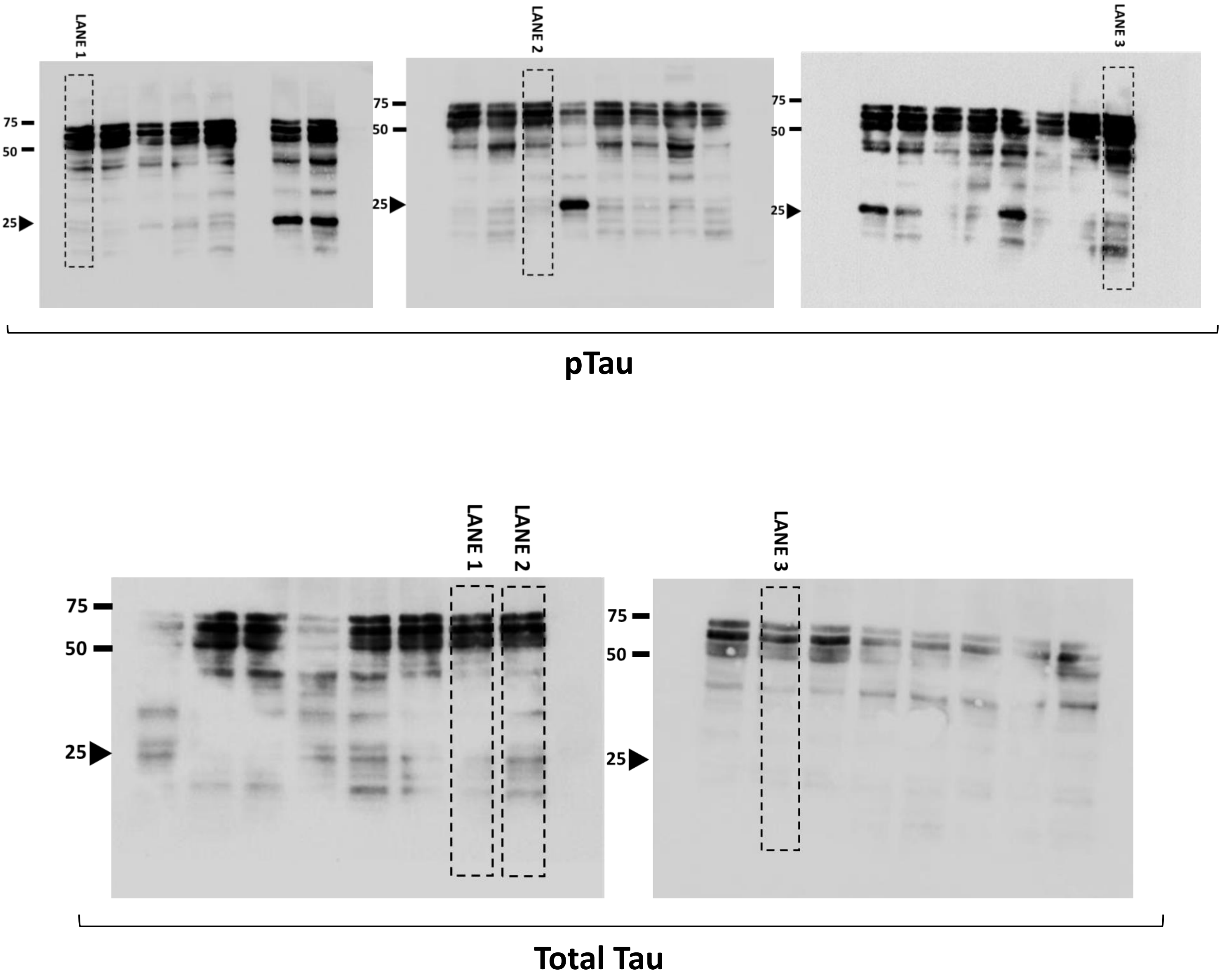

**Figure S5.** Raw data of **Fig.1 – Panel E** western blotting of freeze/thawed rat frontal cortex extracts (lanes used in the main figure are indicated by dotted frames). Asterisks (*) stand for lanes with rat extracts from other cerebral tissue.

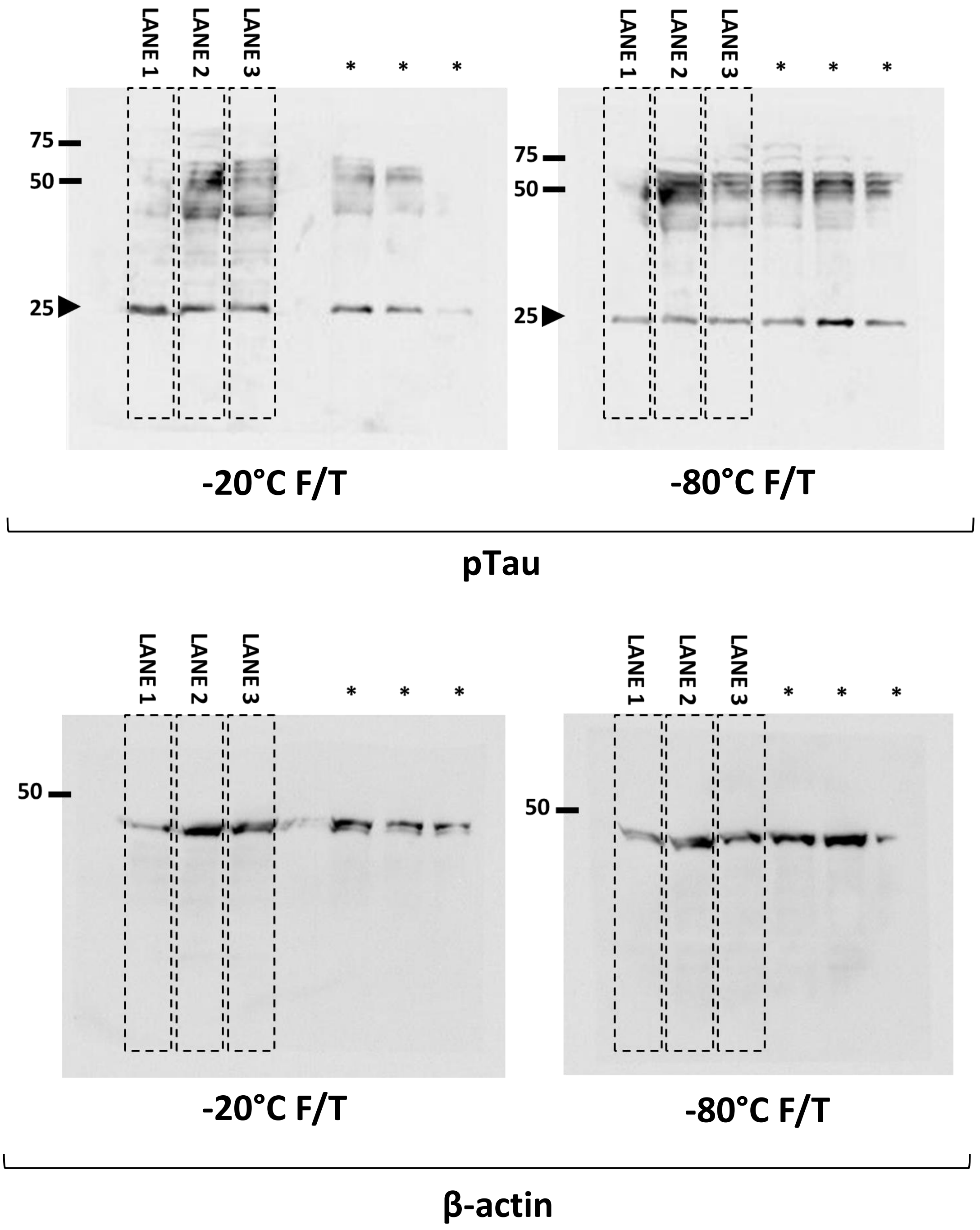

**Figure S6.** Raw data of **Fig.1 – Panel F** western blotting of fresh and freeze/thawed extracts from rat hippocampus and frontal cortex (lanes used in the main figure are indicated by dotted frames).

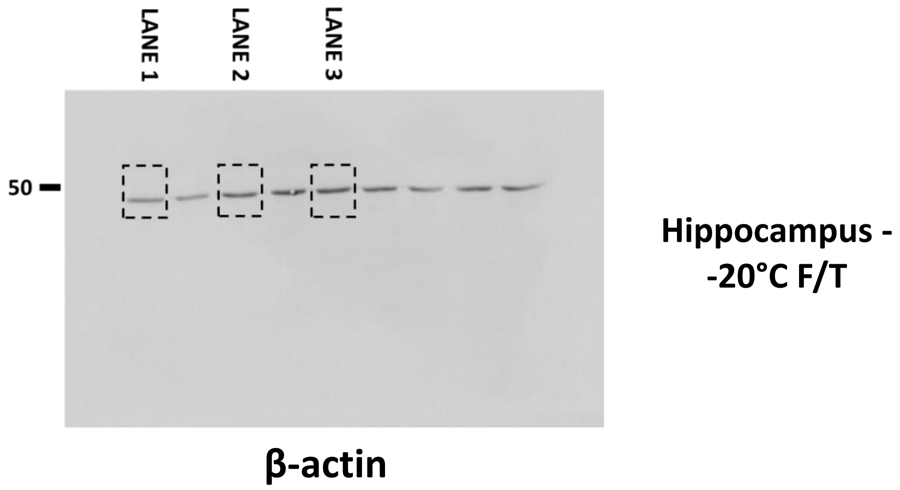


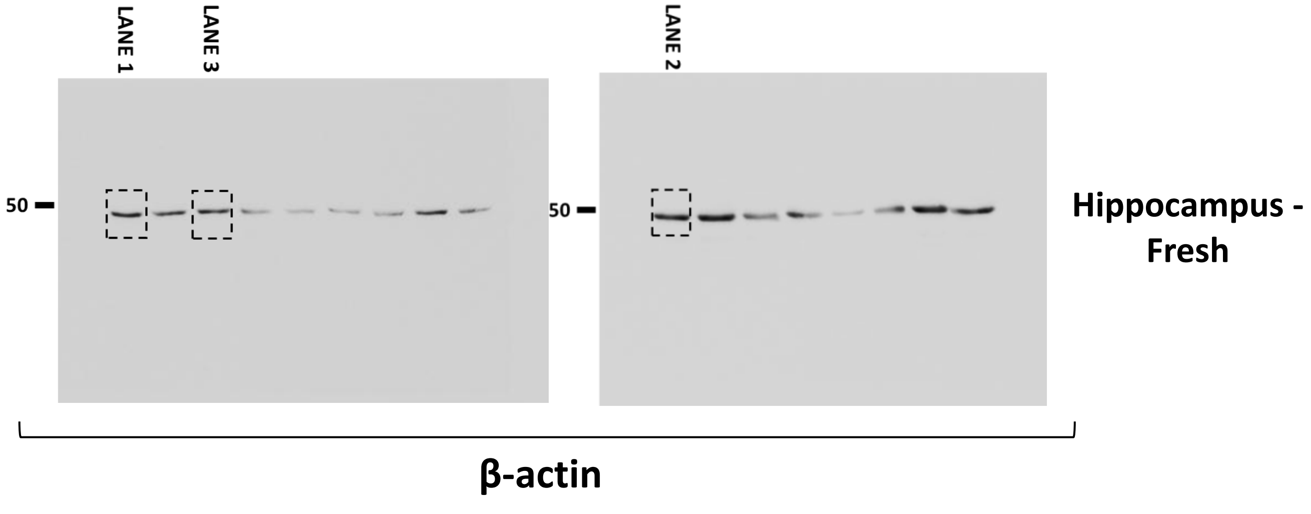

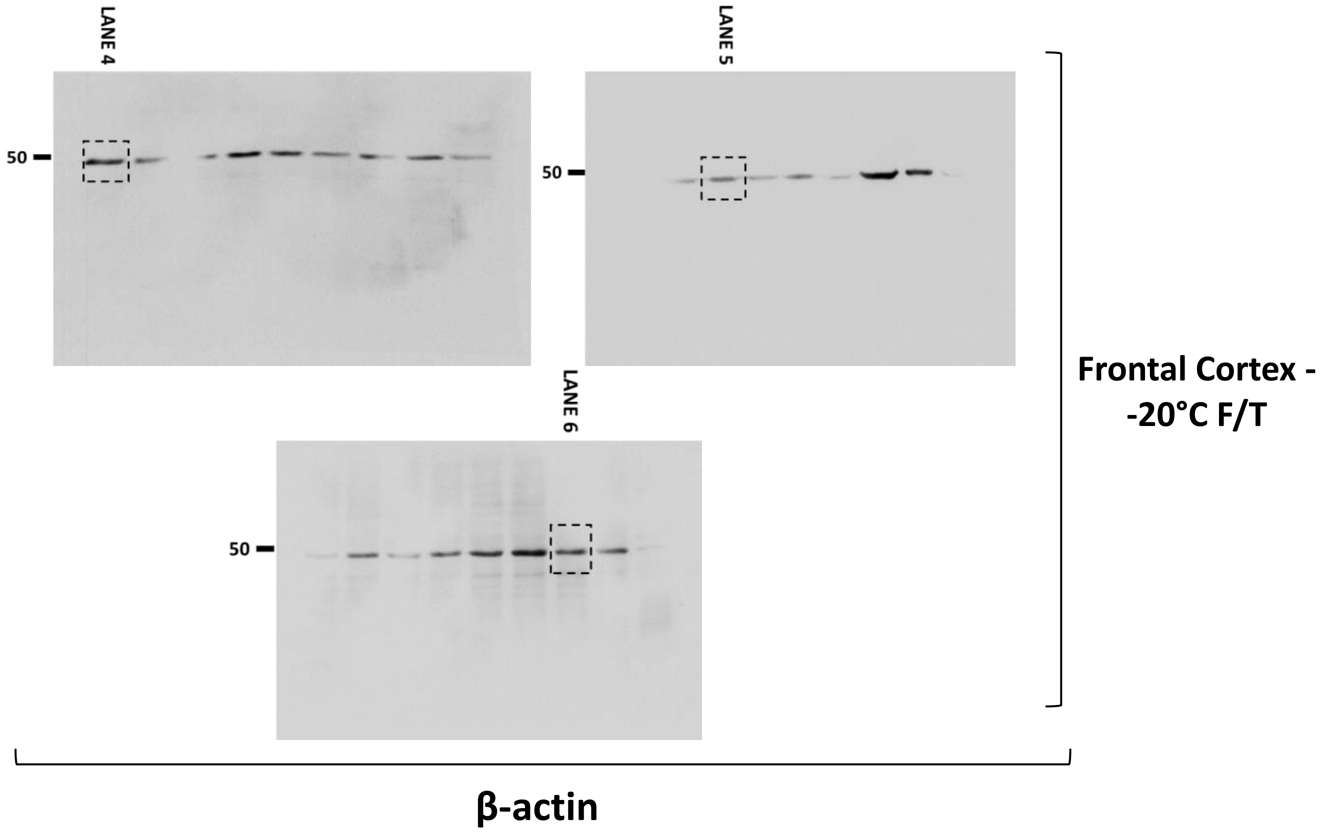
LANE 4
LANE 5
LANE 6
50
50
50
Frontal Cortex -
-20°C F/T
β-actin

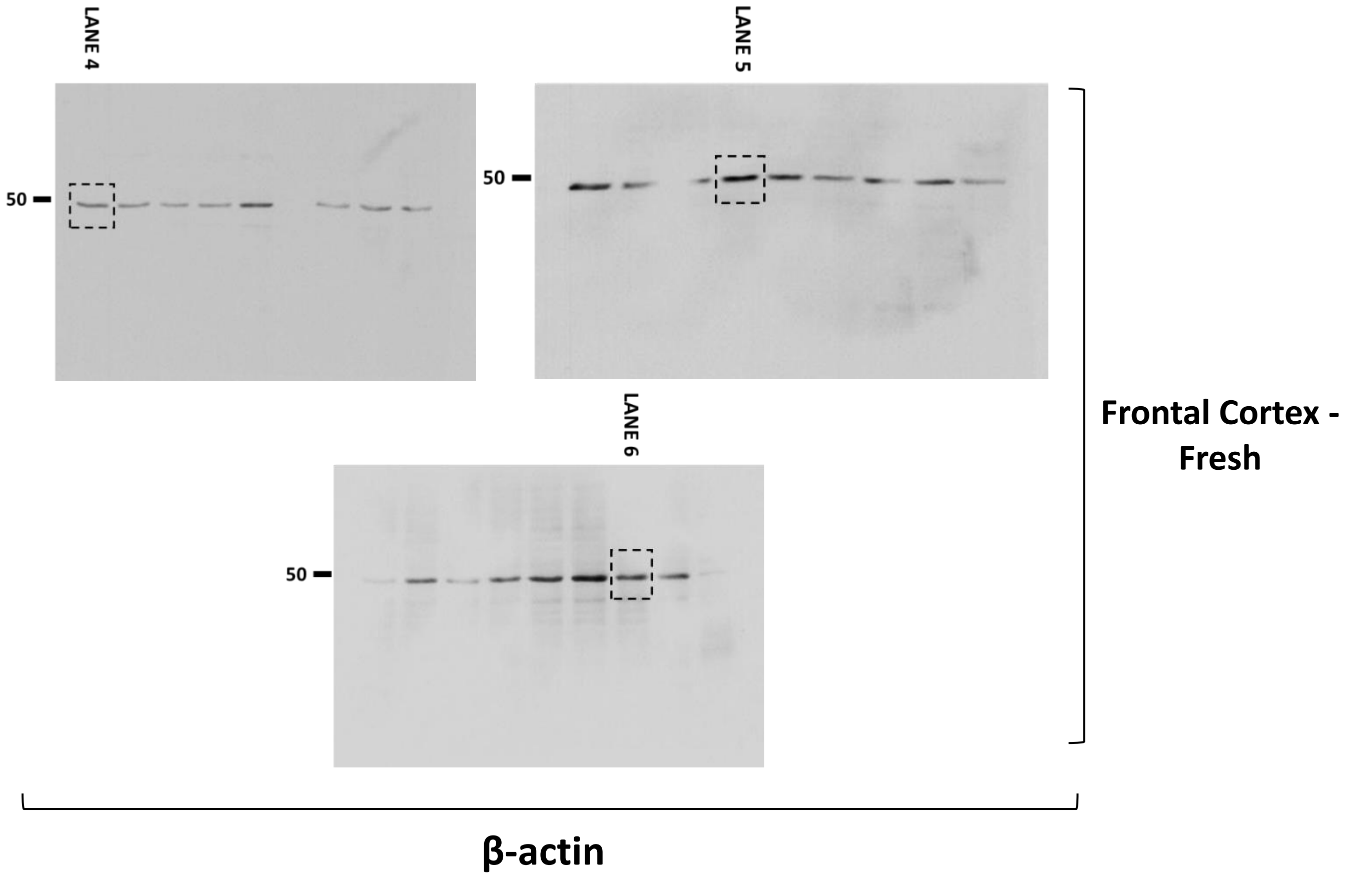
LANE 4
LANE 5
LANE 6
50
50
50
Frontal Cortex -
Fresh
β-actin

**Figure S7.** Raw data of **Fig.3 – Panel A** western blotting of fresh and freeze/thawed extracts from 3xTg mice frontal cortex (lanes used in the main figure are indicated by dotted frames).

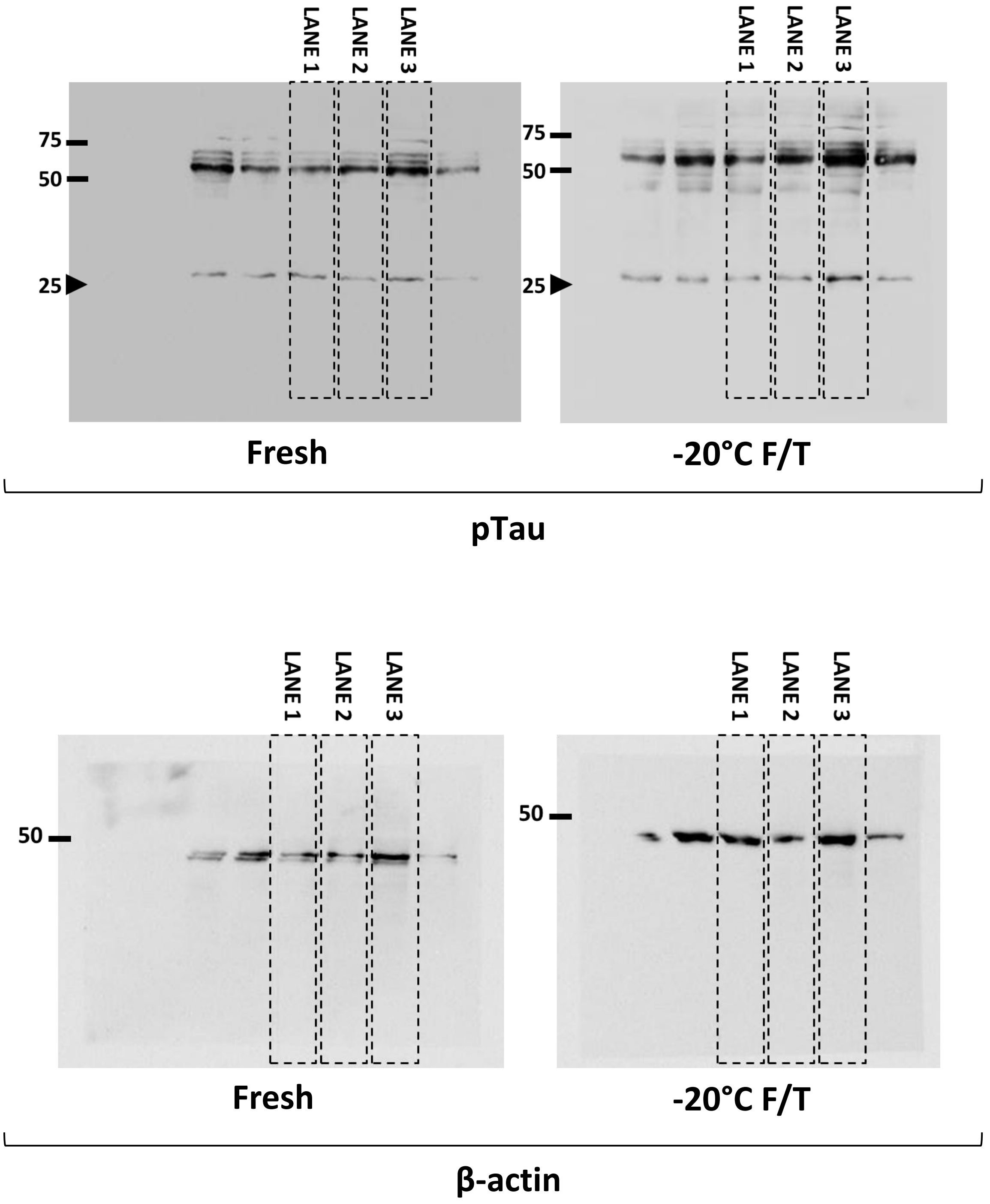

**Figure S8.** Raw data of **Fig.3 – Panel C** western blotting of fresh and freeze/thawed extracts from human frontal cortex (lanes used in the main figure are indicated by dotted frames). Asterisks (*) stand for lanes with human extracts frozen at -80°C.

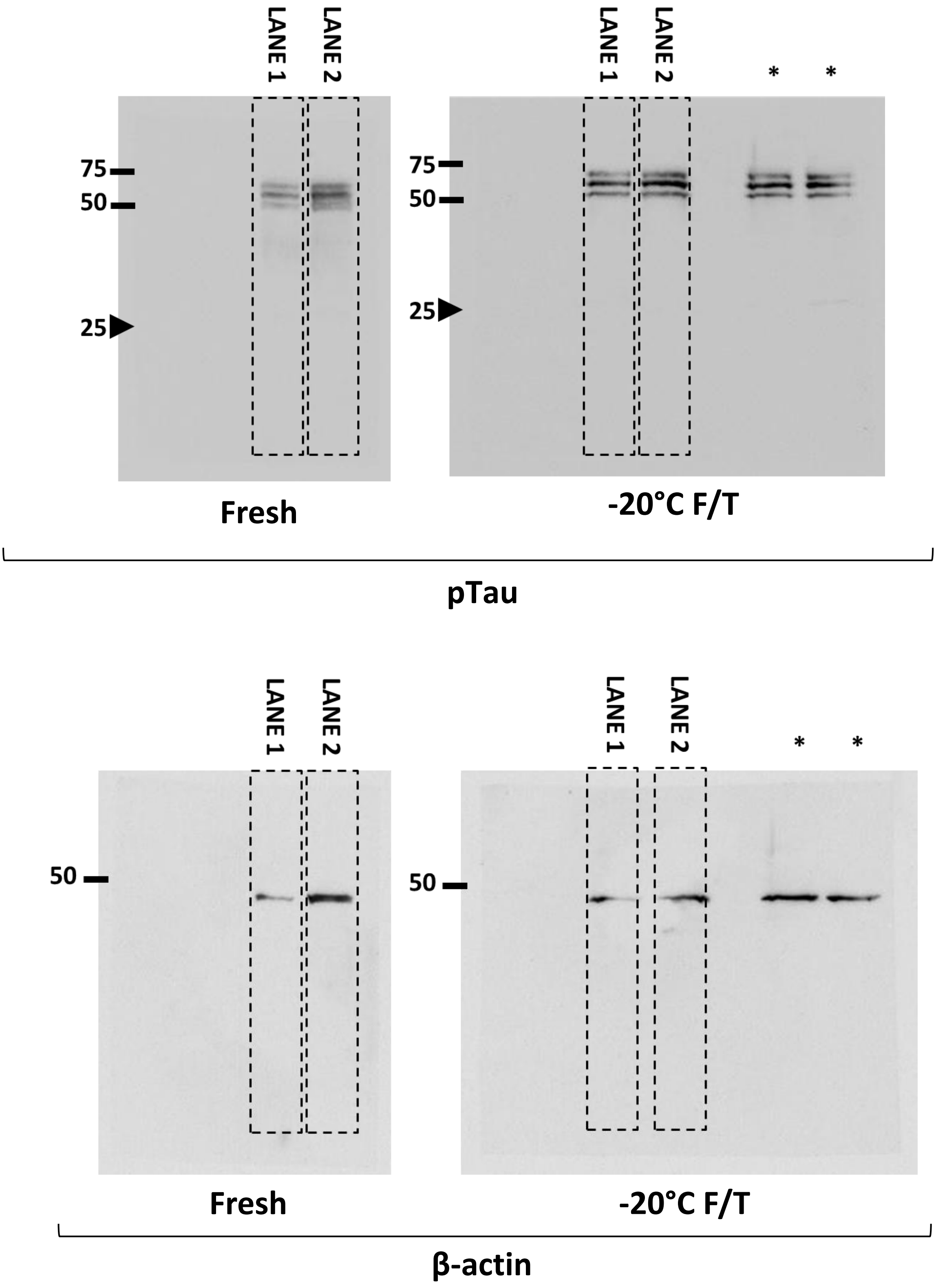
